# Kindlin-3 Mutation in Mesenchymal Stem Cells Results in Enhanced Chondrogenesis

**DOI:** 10.1101/578690

**Authors:** Bethany A. Kerr, Lihong Shi, Alexander H. Jinnah, Jeffrey S. Willey, Donald P. Lennon, Arnold I. Caplan, Tatiana V. Byzova

## Abstract

Identifying patient mutations driving skeletal development disorders has driven our understanding of bone development. Integrin adhesion deficiency disease is caused by a Kindlin-3 (fermitin family member 3) mutation and its inactivation results in bleeding disorders and osteopenia. In this study, we uncover a role for Kindlin-3 in the differentiation of bone marrow mesenchymal stem cells (BMSCs) down the chondrogenic lineage. Kindlin-3 expression increased with chondrogenic differentiation similar to RUNX2. BMSCs isolated from a Kindlin-3 deficient patient expressed chondrocyte markers including SOX9 under basal conditions, which were further enhanced with chondrogenic differentiation. Rescue of integrin activation by a constitutively activated β_3_ integrin construct increased adhesion to multiple extracellular matrices and reduced SOX9 expression to basal levels. Growth plates from mice expressing a mutated Kindlin-3 with the integrin binding site ablated demonstrated alterations in chondrocyte maturation similar to that seen with the human Kindlin-3 deficient BMSCs. These findings suggest that Kindlin-3 expression mirrors RUNX2 during chondrogenesis.

**SUMMARY:** This study by Kerr et al. describes a new role for Kindlin-3 in controlling early chondrocyte differentiation from mesenchymal stem cells and later hypertrophic differentiation of chondrocytes.

## INTRODUCTION

Although our understanding of skeletal development in vertebrates has substantially improved through the use of animal models (Long and Ornitz, 2013), our progress in unraveling human bone development remains limited. Moreover, discrepancies between the effects of inhibitors and genetic ablations in mice (Long and Ornitz, 2013) complicate animal model-based analyses of key pathways underlying endochondral ossification, the main process of skeletal long-bone development. This knowledge is essential not only for understanding development and aging, but also for skeletal disease treatment, orthopaedic injury repair, and bone regenerative medicine. While a number of growth factors and their receptors ranging from FGFs to WNTs as well as transcriptional regulators such as SOX9 and RUNX2 are implicated in bone development, it remains unclear how these signals are integrated within specific bone microenvironments.

Loss-of-function mutations in patients with skeletal development complications represent a unique opportunity to uncover the function of specific proteins in human bone formation. Integrin activation deficiency disease (IADD, also called LAD-III), a rare immunodeficiency, presents in patients as severe bleeding, frequent infections, and osteopetrosis, and is caused by Kindlin-3 (fermitin family member 3) deficiency (Malinin et al., 2009). Kindlins (fermitin family members) function as intracellular adaptors or scaffold proteins involved in inside-out integrin activation by direct binding to the tail of the integrin β subunit (Plow et al., 2009; Moser et al., 2008). A role for Kindlin-2 in chondrogenesis was recently uncovered (Wu et al., 2015). Deletion of Kindlin-2 in mesenchymal stem cells (MSCs) resulted in reduced chondrocyte proliferation and columnar organization. The skeletal defects generated by Kindlin-2 loss indicated a potential role in both intramembranous and endochondral ossification. Further, development of the primary ossification center was impaired leading to limb shortening in mice.

Unlike the ubiquitously expressed Kindlin-2, Kindlin-3 is detected primarily in hematopoietic and endothelial cells. Therefore, the origin of malignant infantile osteopetrosis, i.e., high bone density in Kindlin-3 deficient patients, was traced to impaired bone resorption by osteoclasts originating from the myeloid lineage in bone marrow (Crazzolara et al., 2015; Malinin et al., 2009; Sabnis et al., 2010). However, there is evidence that this osteoclast dysfunction and osteopetrosis is associated with MSC defects (Uckan et al., 2009). The severity of bone problems in Kindlin 3 deficient patients suggests functional defects in other cell types critically involved in bone development and possibly operating in close contact with the hematopoietic stem cell niche. While bone resorption is performed by osteoclasts, differentiation and maturation of MSCs underlie chondrogenesis and ossification. However, there are no reports suggesting either presence or a function of Kindlin-3 in MSCs and bone formation.

In order to understand the mechanisms of bone development in humans we took advantage of bone marrow samples from a previously described Kindlin-3 deficient IADD patients (Malinin et al., 2009) and examined changes in bone marrow-derived mesenchymal stem cells (BMSC) proliferation and adhesion, as well as chondrogenic differentiation. To mechanistically implicate Kindlin-3 as an integrin adaptor, we utilized knock-in mice expressing a Kindlin-3 mutant that is deficient in integrin binding. Utilizing these two models, we demonstrate that Kindlin-3 regulates chondrocyte differentiation and maturation.

## METHODS

### Clinical Samples

Studies were conducted in accordance with the ethical standards of the Institutional Review Board at University Hospitals of Cleveland and with the Helsinki Declaration of 1975, as revised in 2000. Informed consent was obtained from all individuals. Details of the subject’s histories were previously described (Malinin et al., 2009).

### Reagents

All chemicals were obtained from Sigma unless otherwise specified. Media and cell culture reagents were supplied by the Central Cell Services Core of the Lerner Research Institute.

### Bone Marrow Culture

Samples of patient bone marrow before bone marrow transplant were collected (subject). Separately, bone marrow was collected from a normal volunteer (control). Samples were rinsed with Dulbecco’s Modified Eagle’s Medium-low glucose (DMEM-LG) supplemented with 10% FBS (growth media) from a lot selected for expansion and differentiation of human bone marrow-derived mesenchymal stem cells (BMSCs). The marrow cells were then centrifuged on a Percoll gradient as described previously (Solchaga et al., 2004; Lennon and Caplan, 2006). Cells from the top 10 mL of the gradient were collected and seeded into 100 mm tissue culture dishes at a density of 10 × 10^6^ cells per dish; additional dishes were seeded at densities of 5 and 1 × 10^6^ cells for determination of the number of colonies. Cells were cultured until passage four at which point they were plated at 3000 cells per cm^2^ for the following studies. Differentiation of BMSCs along the chondrogenic pathways *in vitro* was completed as described elsewhere (Solchaga et al., 2004; Lennon et al., 1995; Dennis et al., 1998). For chondrogenesis, DMEM-LG was supplemented with 0.1 µM dexamethasone, 1% ITS+Premix, 50 µM ascorbic acid 2-phosphate (Wako), and 1 mM sodium pyruvate (control media). Control medium was then augmented with 10 ng/mL recombinant human TGF-β1 (R&D Systems; chondrogenic media) to induce chondrogenesis. For pellet cultures, BMSCs were placed in 15 mL conical tubes (3 × 10^5^ cells/tube) and centrifuged at 2000 rpm. Cell pellet media was changed to control or chondrogenic after 24 hours.

### Immunofluorescence

Cultures were plated on 10 µg/mL collagen type I coated coverslips in 24 well plates. Cultures were maintained in growth, control, or chondrogenic media, and fixed at 7 days. Fixed cultures were permeabilized by incubation with 0.2% Triton-X100. Cultures were stained with primary rabbit anti-KINDLIN-3 antibody (described previously (Bialkowska et al., 2010)). Cells were then stained with an anti-rabbit Alexa Fluor 488 conjugated secondary antibody (Molecular Probes; RRID: AB_143165) and phalloidin-Alexa Fluor 568 probe (Molecular Probes). Coverslips were mounted onto slides with Vectashield+DAPI (Vector Laboratories; RRID: AB_2336790). Photographs were taken with either a TCS-SPE (Leica; RRID: SCR_002140) or ZMZ1000 (Nikon) microscope with a 20x objective. The mean fluorescence intensity of individual channels was measured using NIH ImageJ (RRID: SCR_003070).

### RNA Isolation and qPCR

RNA was isolated from BMSCs using the Qiagen RNeasy Mini Kit and cDNAs were prepared using the Clontech Advantage RT-for-PCR kit followed by real-time qPCR using the BioRad iQ SYBR green Supermix and measured on a Bio-Rad myIQ2 iCycler. Primers are shown in Table 1. The Kindlin primers were previously described (Bialkowska et al., 2010). Data were analyzed by the ΔΔC_T_ method.

**Table 1.**
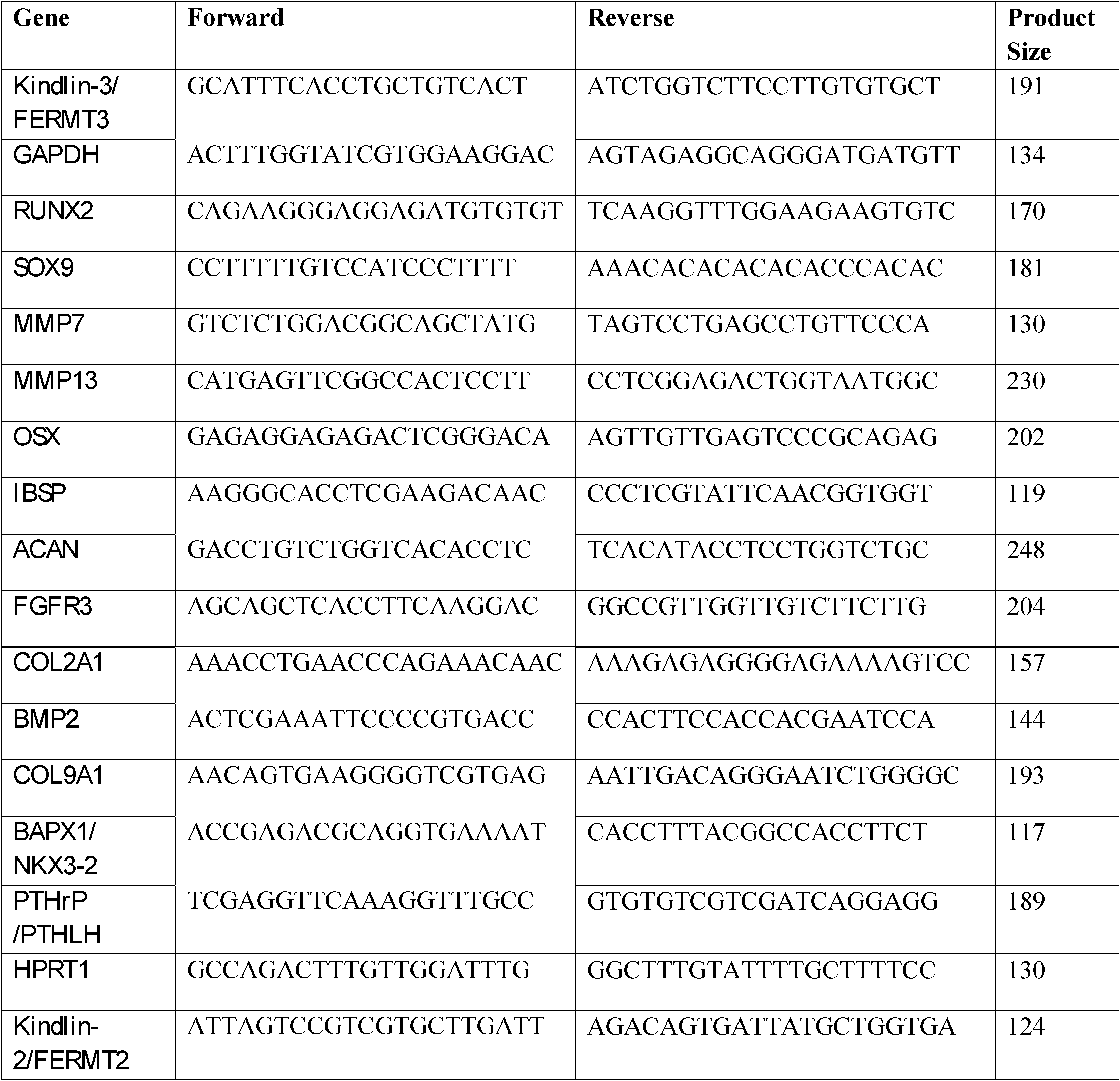
Primers for qPCR.

### Immunoblotting

Control BMSCs were lysed by incubation with RIPA buffer (Pierce) and 1 mM protease inhibitors (Complete Mini, Roche). Samples were incubated with 5X Lane Marker Reducing sample buffer (Pierce) and boiled at 95°C for 10 min. Samples underwent electrophoresis on an 8% polyacrylamide gel at 100 V for approximately 2 hours. Proteins were transferred to a nitrocellulose membrane at 100 V for 1 hour at 4°C. Membranes were blocked in 5% BSA in TBS containing 0.1 % Tween-20. KINDLIN-3 antibody (described previously (Bialkowska et al., 2010)) followed by an IRDye 800CW-conjugated anti-rabbit secondary antibody (LI-COR Biosciences; RRID: AB_621848) was used to visualize proteins. Membranes were scanned with the Odyssey LI-COR infrared scanner in the Lerner Research Institute Imaging Core with excitation at 778 nm and emission at 795 nm.

### Flow Cytometry

Cultures were maintained in growth, control, or chondrogenic media. Cultures were trypsinized at 6, 12, or 23 days, washed, and prepared for flow cytometry. In brief, cells were incubated with human Fc fragment for 15 minutes to block non-specific binding. Cells were stained with directly conjugated antibodies against human proteins: alkaline phosphatase-APC (R&D Systems; RRID: AB_357039), CD73-PE (BD Biosciences; RRID: AB_2033967), CD45-FITC (BD Biosciences; RRID: AB_2732010), CXCR4-PE (R&D Systems), CD146-PE (R&D Systems), CD29-PE (BioLegend; RRID: AB_314320), CD90-FITC (BioLegend; RRID: AB_893429), CD105-FITC (BioLegend; RRID: AB_755956), CD44-FITC (BioLegend; RRID:AB_312957), CD117-PE/Cy7 (BioLegend; RRID: AB_893228), CD11b-PE (eBiosciences; RRID: AB_2043799), CD34-FITC (BioLegend; RRID: AB_1731852), CD133-APC (Miltenyi Biotech). After staining, cells were washed twice in 1X PBS and resuspended in 1% formalin. For intracellular staining, samples were fixed for 10 minutes in 10% formalin. Cells were then stained with anti-human osteocalcin-PE (R&D Systems) in SAP buffer (HBSS with 0.1% saponin and 0.05% sodium azide). Cells were then washed twice with SAP buffer and resuspended in PBS. The stained cells were then analyzed using a BD FACS Canto II running the FACS Diva software. Fluorescence values of stained cells were normalized to a corresponding isotype-fluorochrome labeled control sample.

### Adhesion Assay

Wells of 96 well plates were coated with 1% gelatin, 10 µg/mL collagen type I, 10µg/mL collagen type II (BD Biosciences), 10 µg/mL fibronectin (BD Biosciences), 10 µg/ mL fibrinogen (gift from Dr. Edward Plow), 10 µg/mL vitronectin (R&D Systems), 10 µg/mL laminin (BD Biosciences), 1 mg/mL fibronectin or 1 mg/ mL fibrinogen at 37°C overnight and then blocked with 0.1% BSA for 1 hour prior to plating. We used BSA coated wells as a control for weak substrate attachment. Before plating cells were incubated in a calcein-AM green solution (Invitrogen) for 30 minutes at 37°C. Cells were allowed to attach for one hour. Calcein fluorescence was read at 485 nm excitation and 538 nm emission (SpectraMAX Gemini XS, Molecular Devices). Values were normalized to the BSA control.

### Proliferation Assay

Cells were plated in growth media at a density of 2500 cells/well on either tissue culture plastic or plates coated with 10 µg/mL collagen type I, 10 µg/mL collagen type II, 10 µg/mL fibronectin or 10 µg/mL vitronectin. The CyQUANT NF Cell Proliferation Assay Kit (Molecular Probes) was used to quantify cell numbers after 4 hours, 1, 3, and 5 days. The assay was performed according to the manufacturer’s protocol, and fluorescence was read at 485 nm excitation and 538 nm emission (SpectraMAX Gemini XS, Molecular Devices).

### Histochemical Staining

Matrix produced by cells was assessed histochemically as described previously (Kerr et al., 2008). In brief, sulfated proteoglycans were fixed in 10% formalin and stained for 30 minutes with 1% Alcian Blue 8GS solution (Electron Microscopy Sciences) on day 12 after plating. Alcian Blue staining was quantified by incubation of 4 M guanidine HCl for 15 min. The absorbance of the solution was read at 590nm with a visible plate reader (Vmax, Molecular Devices). Alkaline phosphatase production was assessed histochemically. Cells were incubated with a staining solution of 0.5 mg/mL Naphthol AS-MX phosphate and 1 mg/mL Fast Red in 50 mM Tris-HCl (pH 9.0) for 15 min at 37°C and then fixed with 10% formalin on day 12 after plating. Histochemical staining was quantified by densitometry using NIH ImageJ.

### Biochemical Assays

For alkaline phosphatase activity and DNA content analysis, cultures were lysed in 0.2% Triton X-100 on day 12 after plating. Lysates were incubated with 0.5 mM MgCl_2_, 0.5 mM para-nitrophenol phosphate, and 0.5 M Tris-HCl (pH 9.0) for 30 min at 37°C. The reaction was stopped with 1 N NaOH, and the absorbance was read at 410 nm with a visible plate reader (Vmax, Molecular Devices). A standard curve of alkaline phosphatase activity was created with dilutions of the 4-nitrophenol solution. To normalize samples, DNA content was measured. Lysates were incubated with 0.02 N NaOH and 0.1 g/mL diaminobenzoic acid for 45 min at 65°C. The reaction was stopped with 2 N HCl, and the fluorescence was read at and emission of 420 nm and analysis of 510 nm (SpectraMAX Gemini XS, Molecular Devices). A standard curve was created with calf thymus DNA. Both plate readers use the SOFTmax PRO 4.0 software (Molecular Devices; RRID: SCR_014240).

### Transfection

Subject BMSC’s were plated in 6 well plates (2×10^5^ cells/well) and incubated with 4 μg of WT β_3_ integrin or DR β_3_ integrin DNA per well in the presence of Lipofectamine2000 (McCabe et al., 2007) resulting in a greater than 90% transfection efficiency. After 3 hours, media was changed to control or chondrogenic. After 3 days, cells were lysed for RNA isolation and qPCR as described above.

### Animal Studies

K3KI knock-in mice were generated as previously described (Xu et al., 2014; Meller et al., 2017). Bones were collected from male and female K3KI mice and wild-type (WT) littermates at 9 weeks of age under a Cleveland Clinic IACUC approved animal protocol (*n*=14-17). Bones were fixed in 10% formalin, decalcified in 14% neutral EDTA, and embedded in paraffin. Sections were stained with hematoxylin and eosin (H&E), Toluidine blue, or Safranin O and Fast Green. TRAP (tartrate-resistant acid phosphatase) staining was completed as previously described (McCabe et al., 2011). Immunohistochemistry of bone sections was performed using antibodies against SOX9 (Abcam; RRID: AB_2728660), collagen type X (Abcam; RRID: AB_879742), and collagen type II (Abcam; RRID: AB_731688) after antigen retrieval and hydrogen peroxide treatment. Staining was visualized with DAB. Slides were scanned using a Hamamatsu NanoZoomer by the Virtual Microscopy Core in the Wake Forest School of Medicine. Bone histomorphometry, osteoclast numbers, and growth plate organization were analyzed with the BioQuant Osteo software (RRID: SCR_016423). Immunostaining was quantified using the VisioPharm digital pathology analysis software. Briefly, a region of interest was drawn around the growth plate, and the total cell number within the region counted. We then used custom-designed apps to count the number of positively stained cells within the region.

### Statistical Analysis

Student’s *t* test analysis or one-way ANOVA analysis with Tukey post-test were used to determine statistical significance using GraphPad Prism 7.0 software (RRID: SCR_002798). * represents p<0.05, ** represents p<0.01, *** represents p<0.005.

## RESULTS

### Kindlin-3 is expressed in BMSCs

We identified a point mutation in the Kindlin-3 *FERMT3* gene ablating integrin activation in humans. These patients presented with severe bleeding, frequent infections, and osteopetrosis localized to the proximal area of the growth plate. Bone marrow transplantation resolved the clinical problems (Malinin et al., 2009). Bone marrow-derived mesenchymal stem cells (BMSCs) were isolated from the subject before bone marrow transplantation. To characterize the BMSCs, we assessed established mesenchymal stem cell and hematopoietic stem cell surface markers by flow cytometry (Dominici et al., 2006). Normal and subject BMSC expressed markers of mesenchymal cells: CD90, CXCR4, CD73, CD105, CD146, CD44, and CD29 (Figure S1). Correspondingly, no makers for hematopoietic stem cells were expressed (Figure S1). These verified BMSCs were used in subsequent experiments comparing normal and Kindlin-3 deficient BMSCs.

Kindlin-3 expression was previously demonstrated in cartilage (Ussar et al., 2006), however, its presence in BMSCs remained to be determined. By immunoblotting, normal BMSCs demonstrated Kindlin-3 protein levels equivalent to human umbilical vein endothelial cells (Figure S2A), which were previously shown to express low levels of Kindlin-3 (Bialkowska et al., 2010). To assess how integrin activation might alter Kindlin-3 expression, cells were plated on tissue culture plastic (control), fibronectin, collagen type I, or collagen type II coated dishes. Gene expression of *Kindlin-3/FERMT3* was not altered with culture on different substrates (Figure S2B). To examine how 3D, micromass culture and cell-cell contact might alter Kindlin gene expression, normal BMCS were cultured in a monolayer or in pellets. *Kindlin-3/FERMT3* gene expression was increased by ∼32-fold when cells were grown in 3D pellet cultures (Figure S2C). Conversely, levels of *Kindlin-2/FERMT2* expression were not changed. Thus, Kindlin-3 is expressed in normal BMSCs, and this expression is enhanced under more physiologic, 3D conditions.

### Kindlin-3 is induced during chondrogenesis

As the 3D pellet is representative of the mesenchymal condensation during chondrogenesis (Solursh et al., 1982), we closely examined Kindlin-3 expression in normal BMSCs on gelatin-coated coverslips after chondrogenic induction. Stimulation of chondrogenesis induced cytoplasmic Kindlin-3 protein expression 9.6-fold in cultured BMSCs (Figure 1A). In addition, actin filament formation was increased 6.5-fold in cells treated with chondrogenic media compared with BMSCs cultured in control media (Figure 1A). Consistent with protein expression, gene expression of *Kindlin-3/FERMT3* increased with time in chondrogenic media, showing its highest expression at day 14 when it was 5.3-fold higher than BMSCs cultured in control media (Figure 1B). Thus, Kindlin-3 expression mirrored the increase over time of the terminal chondrogenic marker *RUNX2* (runt-related transcription factor 2) expression (Figure 1C) (Chen et al., 2014; Inada et al., 1999). Thus, Kindlin-3 expression parallels RUNX2 expression during chondrogenesis in BMSC.

**Figure 1.**
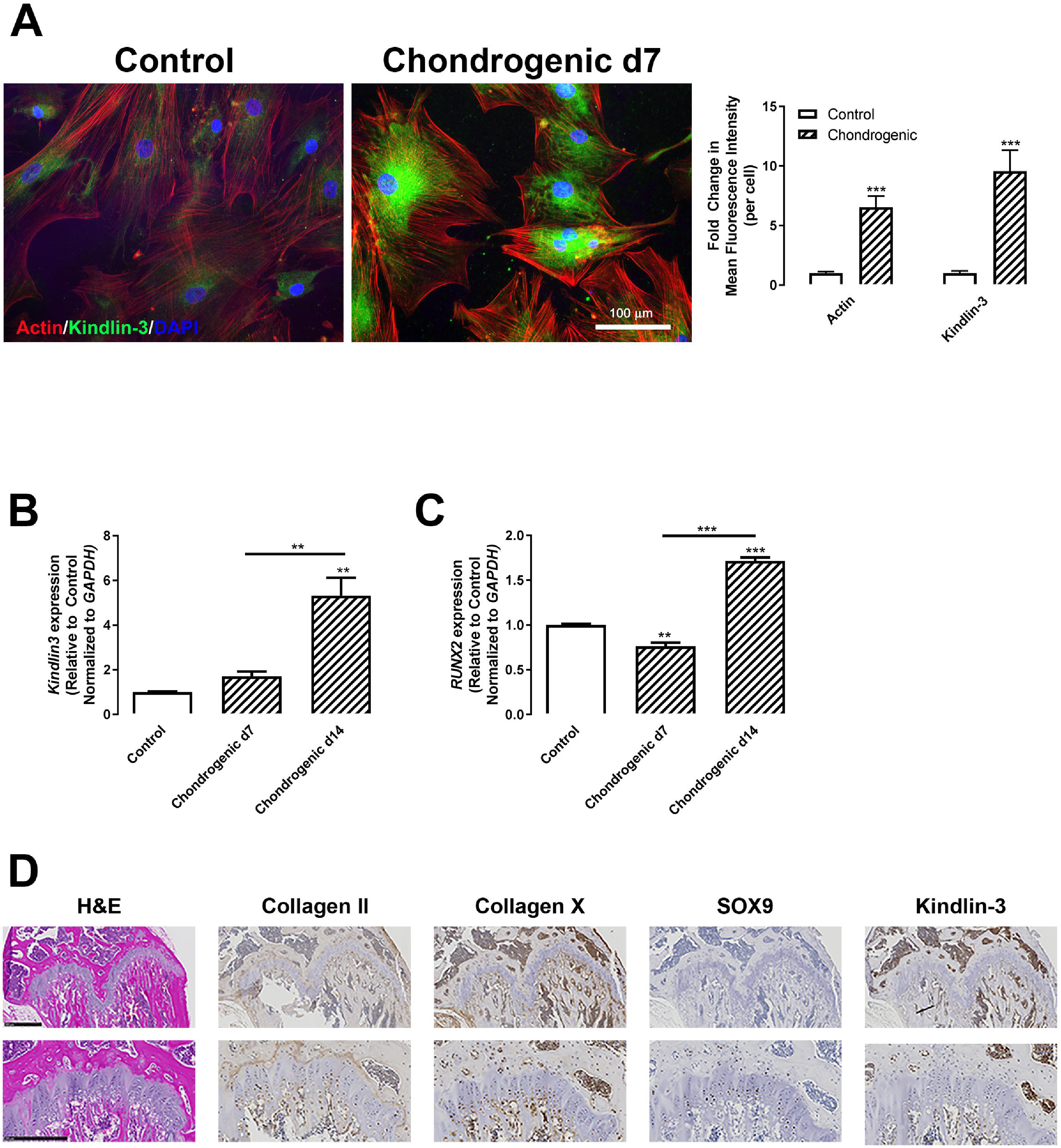
Kindlin-3 is upregulated with chondrogenic differentiation in BMSCs and growth plates. BMSC isolated from normal patients were treated with control (open bars) or chondrogenic (patterned bars) media for 7 (A-C) or 14 days (B-C). (A) Cells were grown on 10 μg/mL collagen type I coated glass coverslips were permeabilized and stained for actin (red), Kindlin-3 (green) or nuclei (blue). Mean fluorescence intensity of actin and Kindlin-3 were measured and represented as mean per cell±SEM. (B-C) Gene expression of *Kindlin-3* (B) or *RUNX2* (C) normalized to *GAPDH* represented as mean fold change from control media±SEM. (D) Sections of 9-week WT mice bones stained for hematoxylin & eosin (H&E), Collagen type II, Collagen type X, SOX9, and Kindlin-3. Scale bars represent 500 μm (top) and 250 μm (bottom). ** represents p<0.01 and *** represents p<0.005 by Student’s *t* test (A) or one-way ANOVA (B-C).

To ascertain Kindlin-3’s role in chondrocyte maturation, 9-week old WT murine growth plates were stained for Kindlin-3 and markers of chondrocyte differentiation (Figure 1D). Collagen type II and SOX9 were localized to proliferative and columnar chondrocytes, while collagen type X was found predominately in hypertrophic chondrocytes (Figure 1D). Kindlin-3 was highly expressed in bone marrow cells in accordance with prior studies demonstrating its presence in hematopoietic stem cells and immune cells. In the growth plate, Kindlin-3 staining was found in both early and hypertrophic chondrocytes (Figure1D), indicating that Kindlin-3 may function at two distinct points during chondrogenesis.

### Kindlin-3 deficiency alters BMSC adhesion, proliferation, and differentiation

In our initial characterization of the Kindlin-3 deficient patient, we demonstrated that a loss of Kindlin-3 diminished lymphocyte adhesion by disabling integrin activation (Malinin et al., 2009). Comparing the adhesion of normal and subject BMSCs to collagen type I, collagen type II, fibronectin, and vitronectin, we confirm that the Kindlin-3 deficient BMSCs displayed decreased attachment (Figure 2A) likely due to the lack of integrin activation. Interestingly, these BMSCs proliferated faster on all substrates tested (Figure 2B-F) demonstrating that the diminished adhesion permits a higher rate of proliferation. This is in line with prior studies demonstrating that focal adhesions dissolve and attachment to the extracellular matrix is reduced during cell division (Jones et al., 2018; Fang et al., 1996; Matus et al., 2014).

**Figure 2.**
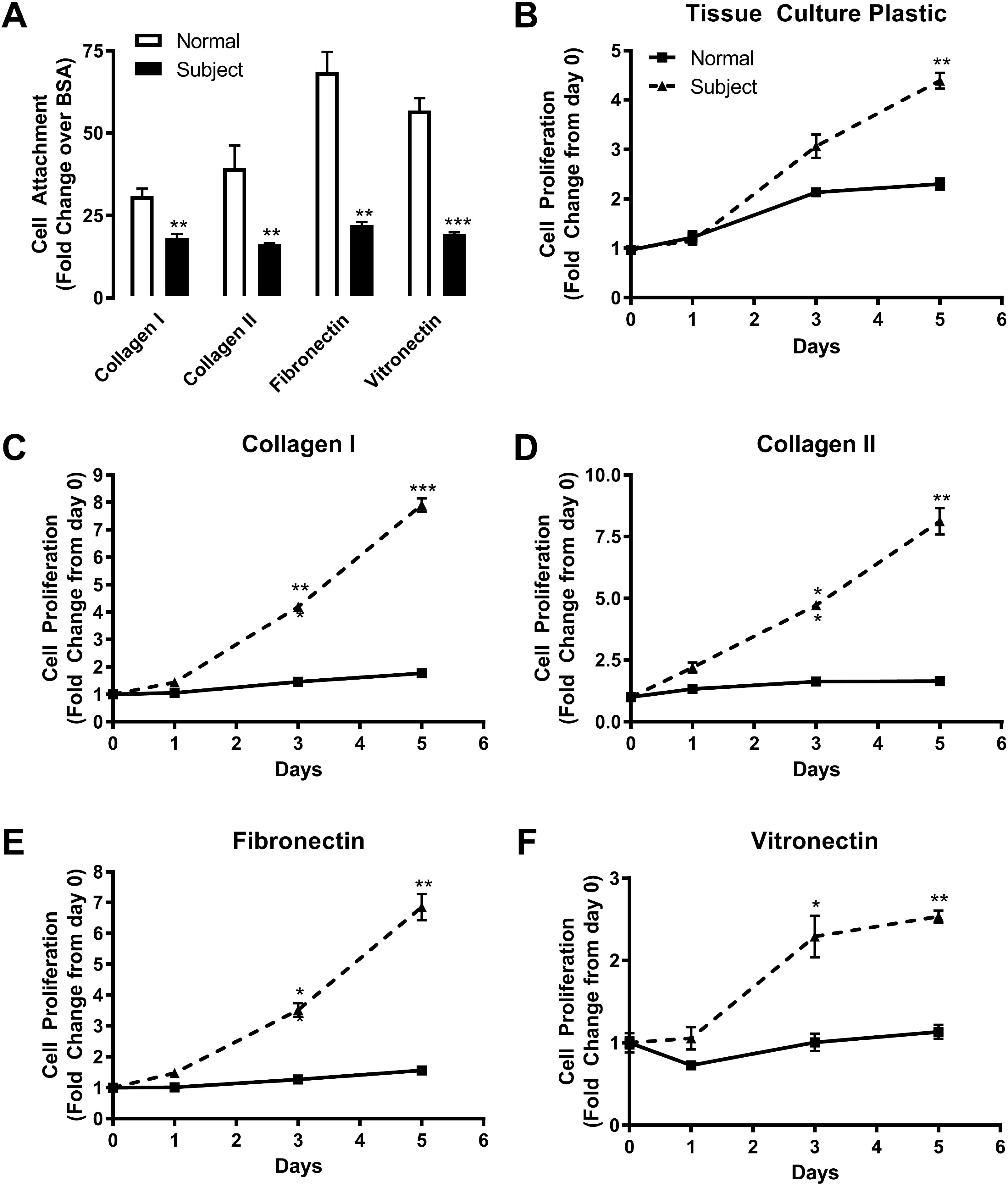
Kindlin-3 deficiency inhibits substrate adhesion but stimulates proliferation. BMSC from normal controls or the Kindlin-3 deficient subject were plated on 10μg/mL collagen type I, 10μg/mL collagen type II, 10μg/mL fibronectin, or 10μg/mL vitronectin with tissue culture plastic coated with BSA as a control. (A) Adherence of cells to the substrates was measured after 30 minutes and represented as fold change from BSA control±SEM. (B-D) The proliferation of BMSC was measured over 5 days and represented as fold change from day 0±SEM. * represents p<0.05, ** represents p<0.01, and *** represents p<0.005 by Student’s *t* test.

Our previous study demonstrated that the subject’s BMSCs stimulated increased differentiation of bone and cartilage compared to control and subject post BMT cells when loaded into ceramic cubes and injected into immunocompromised mice (Malinin et al., 2009). To determine how Kindlin-3 deficiency alters BMSC differentiation, we analyzed normal and subject cells in a variety of chondrogenic assays. First, cells were plated on collagen type I and grown in control media or chondrogenic media containing TGF-β1 and proteoglycan accumulation was measured as a marker for chondrocyte differentiation. Alcian blue staining to visualize proteoglycan deposition and densitometric analysis demonstrated increased proteoglycan release by the subject cells even in control media, while in chondrogenic media, proteoglycan release was further enhanced 2.5-fold in subject Kindlin-3 deficient cells (Figure 3A) Chondrogenesis is also marked by a decrease in alkaline phosphatase production. First, alkaline phosphatase expression was examined histochemically. In control media, subject cells displayed 2-fold decreased alkaline phosphatase expression compared with normal BMSCs (Figure 3B). Second, alkaline phosphatase activity was measured biochemically, and a 5.8-fold decrease was found in the subject BMSCs compared with normal cells in control media (Figure 3C).

**Figure 3.**
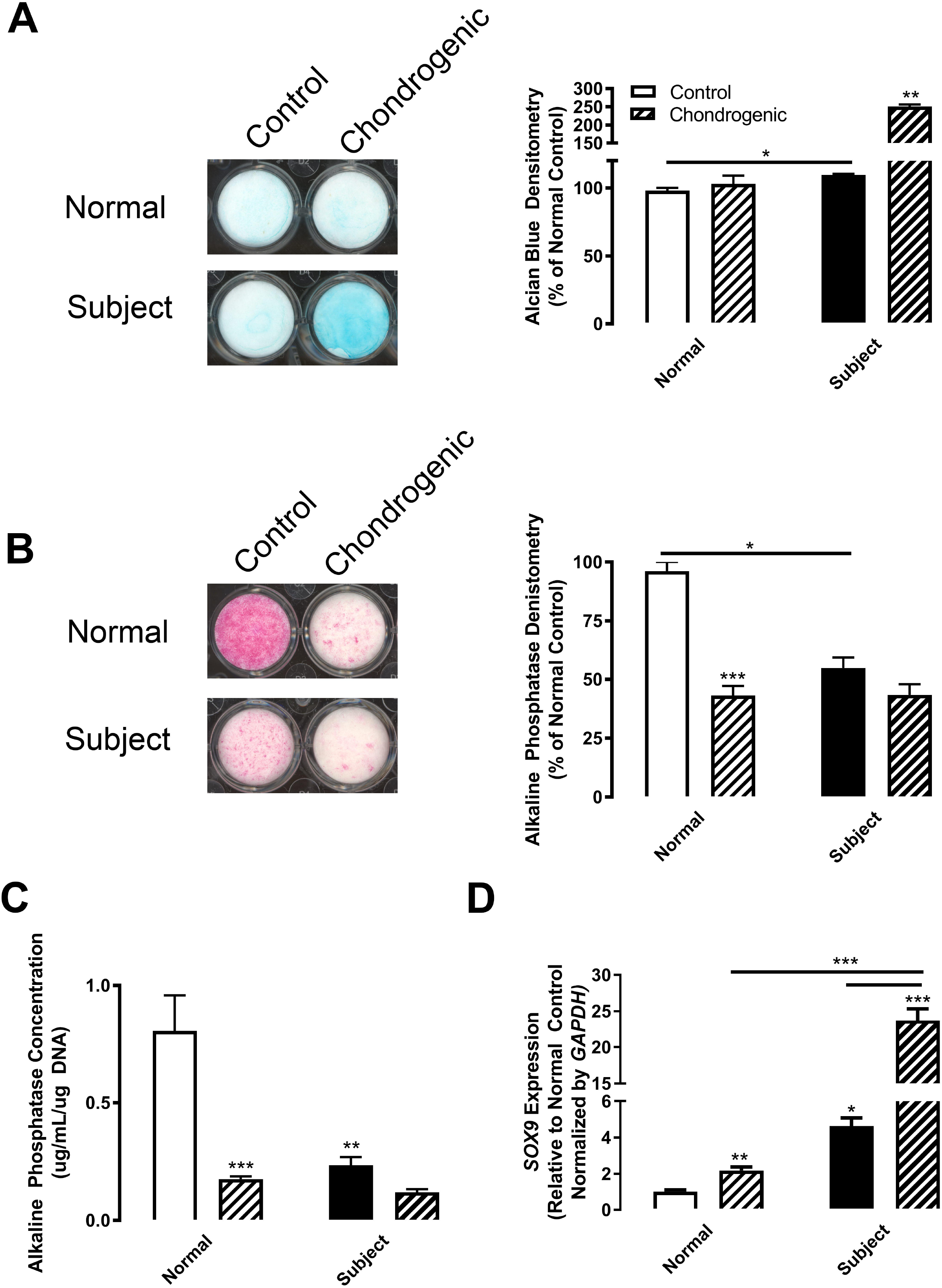
Kindlin-3 deficiency enhances chondrogenesis. BMSC from normal controls (white bars) or the Kindlin-3 deficient subject (black bars) were treated with either control (open bars) or chondrogenic (patterned bars) media. (A) Alcian Blue staining was measured after 12 days and represented as mean percent of control densitometry±SEM. (B) Alkaline Phosphatase staining was measured after 12 days and represented as mean percent of control densitometry±SEM. (C) Alkaline phosphatase activity was assessed biochemically and represented as mean concentration per μg DNA±SEM. (D) Gene expression of *SOX9* normalized to *GAPDH* represented as mean fold change from control media±SEM. * represents p<0.05, ** represents p<0.01, and *** represents p<0.005 by one-way ANOVA.

Finally, *SOX9* expression was examined as a marker of chondrogenic initiation. SOX9 (Sex Determining Region Y-Box 9) is required for chondrogenesis, and its expression inhibits the transition into hypertrophy and terminal differentiation by decreasing RUNX2 expression (Dy et al., 2012). The expression of *SOX9* was 4.6-fold higher in subject Kindlin-3 deficient BMSCs in control media compared with normal cells. *SOX9* expression was upregulated by chondrogenic media (TGF-β1 treatment) as expected, although subject cells had 10.8-fold higher expression compared with normal BMSCs (Figure 3D). These data indicate that even prior to chondrogenesis initiation the Kindlin-3 null BMSCs are already advanced along the chondrogenic differentiation pathway.

To confirm that the differentiation of BMSCs from the Kindlin-3 deficient subject was more advanced, we examined the expression of a variety of chondrogenesis-related genes in BMSCs cultured in control media (Figure 4). Most of the genes associated with chondrocyte maturation (Majumdar et al., 2003) were upregulated in BMSCs isolated from the Kindlin-3 deficient subject with the exception of *RUNX2* expression which was significantly lower in the subject BMSCs (Figure 4) and is required for endochondral ossification and late chondrocyte maturation (Chen et al., 2014; Inada et al., 1999). Genes upregulated in the subject’s BMSC in the absence of differentiation media included matrix metalloproteinases required for chondrocyte hypertrophy: *MMP7* (3.6-fold) and *MMP13* (5.8-fold) (Mackie et al., 2008), and extracellular matrix proteins associated with chondrocyte maturation: collagens (*COL2A1*, 115.8-fold and *COL10A1*, 311.8-fold) (Häusler et al., 2002), aggrecan (*ACAN*, 73.7-fold), and bone sialoprotein/Spp1/osteopontin (*IBSP*, 41.9-fold). Additionally, transcription and growth factor-related genes driving early chondrocyte differentiation were higher in subject BMSCs: Osterix/Sp7 (*OSX*, 14.7-fold), SOX9 (*SOX9*, 8.4-fold), BAPX1/NKX3-2 (*BAPX1*, 981.5-fold), bone morphogenic protein 2 (*BMP2*, 191.5-fold), parathyroid hormone-related protein (*PTHRP*, 1685.1-fold), and fibroblast growth factor receptor 3 (*FGFR3*, 140.2-fold). Thus, the genes associated with early chondrocyte differentiation and SOX9 expression were upregulated in Kindlin-3 deficient BMSCs without any stimulation, while RUNX2 expression was decreased which could prevent terminal differentiation.

**Figure 4.**
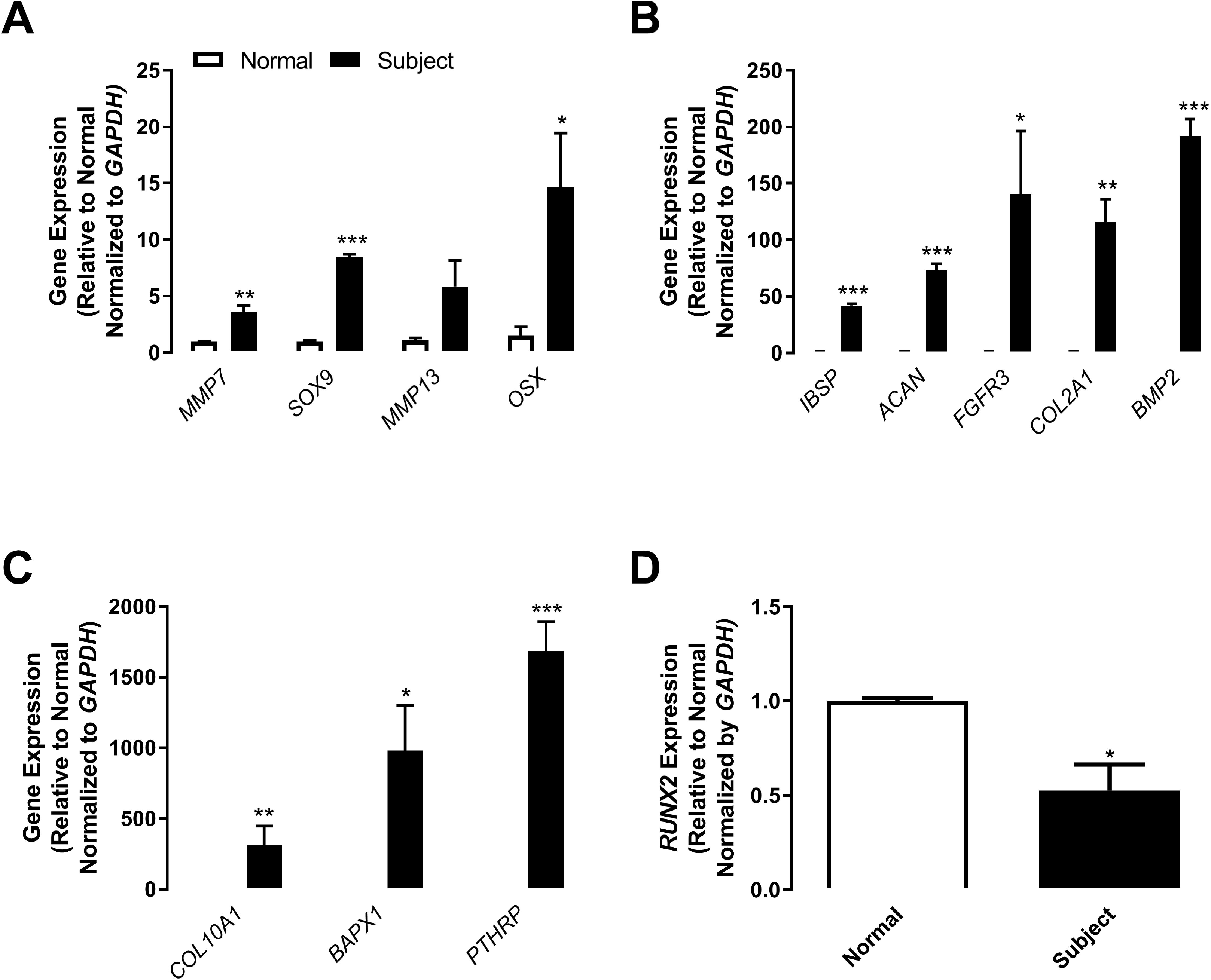
Kindlin-3 deficient BMSCs are further along the pathway towards chondrogenesis. BMSC from normal controls (white bars) or the Kindlin-3 deficient subject (black bars) were maintained in control, growth media. (A-D) Gene expression of the labeled genes was normalized to *GAPDH* represented as mean fold change from control media±SEM. * represents p<0.05, ** represents p<0.01, and *** represents p<0.005 by Student’s *t* test.

### Kindlin-3 deficiency is rescued by constitutive integrin activation

To evaluate whether expression of constitutively active integrin could bypass the requirement for Kindlin-3 and thereby rescue abnormalities caused by Kindlin-3 deficiency, subject BMSCs were transfected with either WT β_3_ integrin or a constitutively active DR β_3_ integrin mutant (McCabe et al., 2007). As anticipated, active integrin expression enhanced subject BMSC adhesion ∼3-fold on collagen type I, collagen type II, and fibronectin substrates demonstrating that expression of active but not WT integrin bypasses the requirement for Kindlin-3 for integrin-mediated adhesion (Figure 5A). Expression of active but not WT integrin was able to revert subject BMSCs to an earlier stage in chondrogenic differentiation as demonstrated by diminished *SOX9* expression in cultures treated with chondrogenic media (Figure 5B). Thus, expression of activated β_3_ integrin overcomes the Kindlin-3 deficiency rescuing cell adhesion and the associated differentiation.

**Figure 5.**
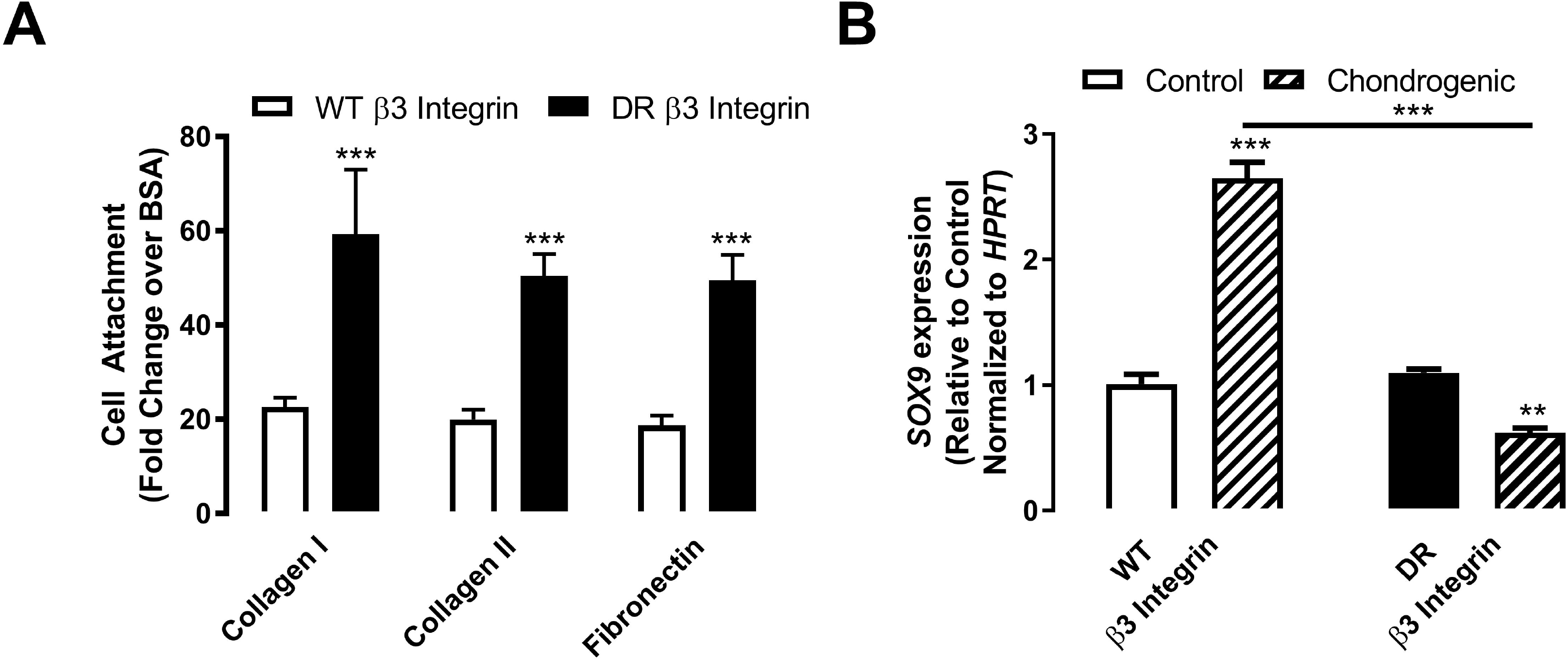
Constitutively active β3 integrin expression rescues Kindlin-3 deficient attachment and chondrogenesis. BMSC from the Kindlin-3 deficient subject were transfected with either control WT β3 integrin (white bars) or constitutively active DR β3 integrin (black bars). (A) Cell attachment was measured as mean as fold change from BSA control±SEM. (B) BMSC were treated with control (open bars) or chondrogenic (patterned bars) media. Gene expression of *SOX9* normalized to *HPRT1* represented as mean fold change from control media±SEM. * represents p<0.05, ** represents p<0.01, and *** represents p<0.005 by Student’s *t* test.

### Mutated Kindlin-3 mice display disrupted growth plate chondrogenesis

To confirm the functional role of Kindlin-3 integrin axis in chondrogenesis, we examined the growth plates of Kindlin-3 mutant mice. The K3KI mutated Kindlin-3 knock-in mice contain a double mutation (Q^597^W^598^/AA) designed to disrupt the integrin recognition region resulting in diminished integrin binding (Xu et al., 2014, 2015). Integrin expression in these mice remains equal to WT mice (Meller et al., 2017). Growth plates in 9-week old K3KI and WT mice were stained for markers of cartilage maturation. The overall structure of the growth plate was examined via Movat’s pentachrome staining, while Safranin O and Fast Green were used to differentiate the growth plate area (Figure 6A). The trabecular bone volume appears higher in K3KI mice, while growth plates between WT and K3KI mice are approximately the same size. Mutation of Kindlin-3 resulted in the continued presence of hypertrophic cells in the ossification front of K3KI bones, while hypertrophic chondrocytes underwent terminal differentiation and were lost in the WT bones (Figure 6A). The expression and localization of the chondrocyte differentiation markers SOX9, collagen type II, and collagen type X were visualized by immunohistochemistry (Figure 6B). All three makers demonstrated 1.2-fold higher staining in the K3KI growth plates compared with WT growth plates confirming that the integrin-binding function of Kindlin-3 drives the changes in chondrogenic maturation markers seen in BMSCs. Additionally, bone histomorphometry of H&E and TRAP stained bones demonstrated 2.3-fold increased bone formation (BV/TV) with an associated 1.3-fold decrease in trabecular spacing but no change in trabecular number (Figure S3A-C). These data are in accordance with the osteopetrosis seen in Kindlin-3 deficient patients (Yarali et al., 2003; Malinin et al., 2009). Further, TRAP staining demonstrated increased numbers of osteoclasts in K3KI mice (Figure S3D-E) similar to Kindlin-3 deficient mice (Schmidt et al., 2011). These changes in the growth plate structure indicate a second potential mechanism for the osteopetrosis in Kindlin-3 deficient, IADD patients.

**Figure 6.**
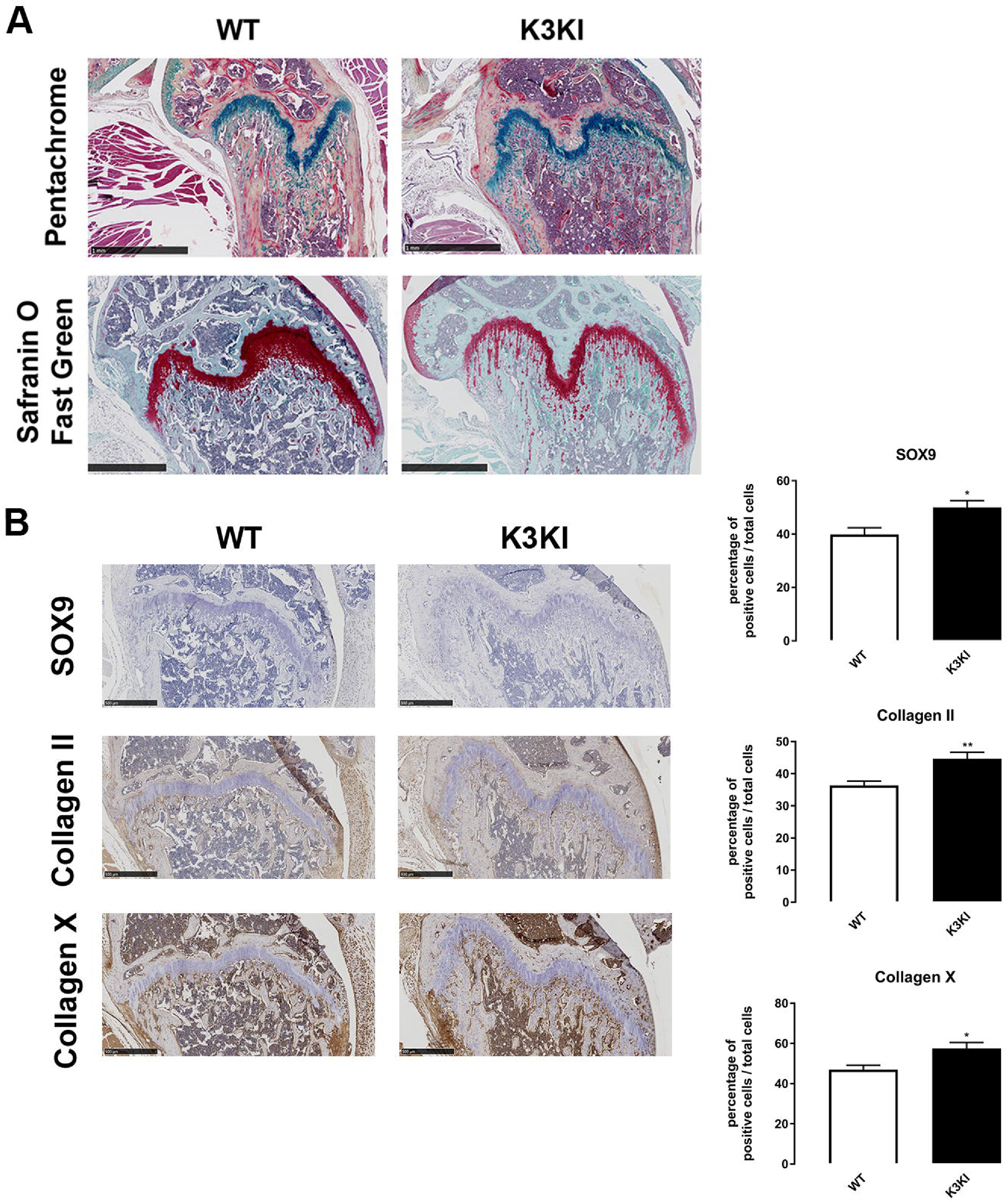
Blocking Kindlin-3 integrin binding results in growth plate abnormalities. Hindlimbs from 9-week old K3KI mutant and wild-type (WT) mice were sectioned and stained with (A) pentachrome or safranin O counterstained with fast green to visualize the growth plate area. Scale bars represent 1 mm. (B) Hindlimbs were also immunostained for SOX9, Collagen type II, and Collagen type X and the numbers of cells stained per total cells in the growth plate calculated as mean±SEM. * represents p<0.05 and ** represents p<0.01 by Student’s *t* test.

## DISCUSSION

In this study, we demonstrate that Kindlin-3 regulates chondrogenesis in MSCs. Temporal Kindlin-3 expression was associated with BMSC differentiation towards the chondrocyte lineage and with increased chondrocyte maturation. As with cells of the hematopoietic lineage, Kindlin-3 loss diminished BMSC adhesion to a variety of substrates resulting in increased proliferation. Under basal conditions, BMSCs lacking Kindlin-3 expressed chondrocyte markers which were further upregulated upon chondrogenic induction. Constitutive integrin activation in Kindlin-3 deficient BMSCs rescued cell adhesion and reduced *SOX9* expression. Growth plates in mice with Kindlin-3 integrin binding blocked demonstrate disorganized growth plates with increased late chondrocyte differentiation and retention of hypertrophic chondrocytes in the trabeculae. Taken together, our data suggest that Kindlin-3 expression is increased in a biphasic fashion during chondrocyte differentiation in a manner similar to RUNX2 (Figure 7).

**Figure 7.**
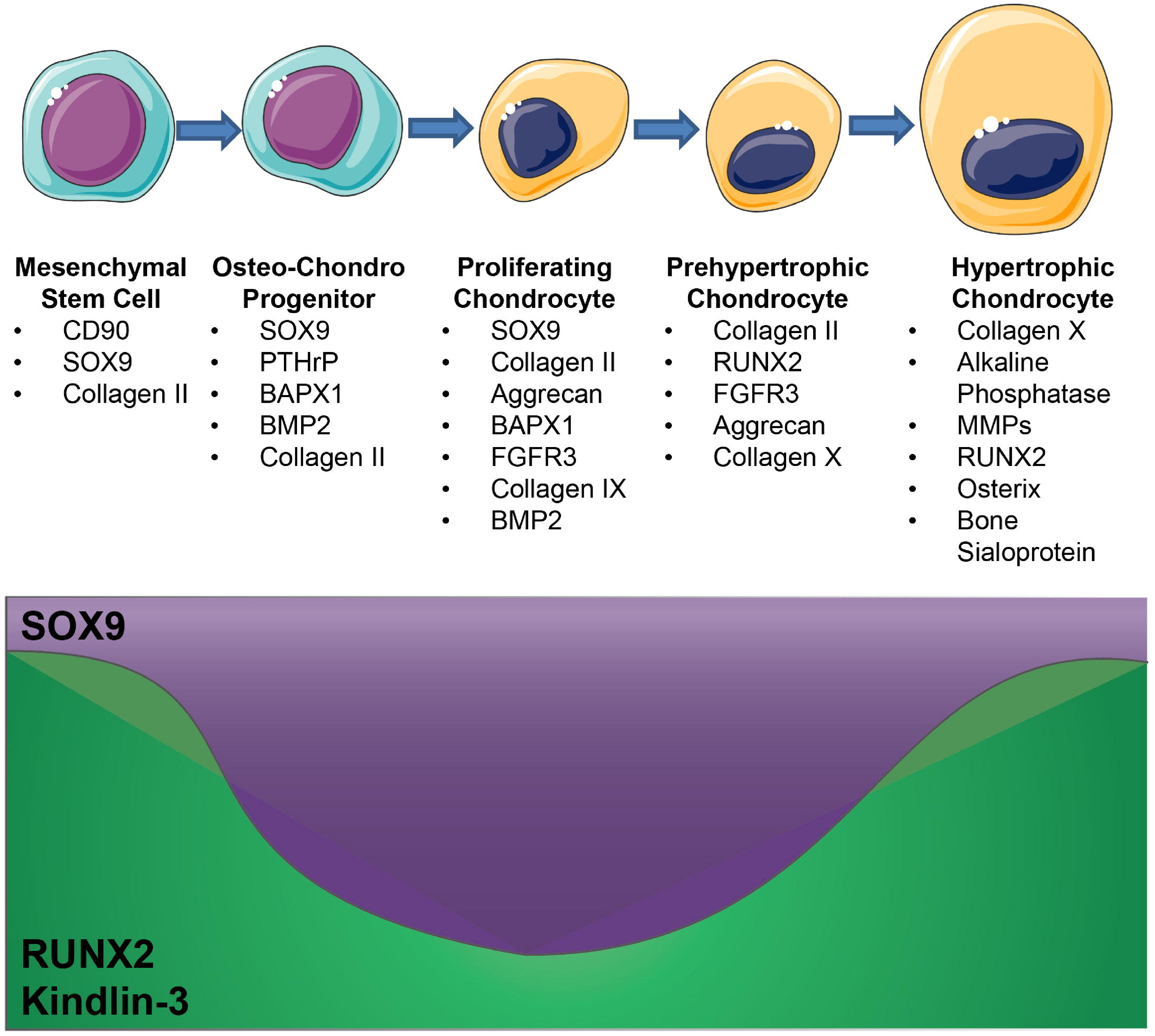
Kindlin-3 Expression during chondrogenesis. As BMSCs differentiate into hypertrophic chondrocytes changes in SOX9 and RUNX2 expression drive changes in differentiation marker expression. The effects of Kindlin-3 deficiency closely resemble a decrease in RUNX2 expression and, thus Kindlin-3 likely has biphasic expression during chondrogenesis.

### Kindlin-3 may play a role in the balance between RUNX2 and SOX9

RUNX2 is the main transcription factor regulating chondrocyte hypertrophy and terminal differentiation (Chen et al., 2014; Inada et al., 1999). Our data demonstrate that Kindlin-3 expression increases in MSCs over time when exposed to chondrogenic media paralleling the increase in RUNX2 in culture. Additionally, in growth plates, Kindlin-3 was expressed early in resting chondrocytes and again in hypertrophic chondrocytes just prior to the ossification front. This pattern of expression is similar to RUNX2. Loss of Kindlin-3 allowed for initial chondrocyte differentiation of BMSCs and increased SOX9 production. Similarly, in mice with RUNX2 deficiencies, SOX9 and parathyroid hormone-related protein levels were equal to that of control mice, and early chondrogenesis proceeded normally (Amizuka et al., 1994). In fact, one study demonstrated that premature stimulation of chondrocyte hypertrophy in MSCs resulted in enhanced calcification and bone formation (Pelttari et al., 2006). This presents one potential mechanism for why the loss of Kindlin-3 induces osteopenia in patients. Since the BMSCs derived from patients had premature stimulation of chondrogenesis, they could induce excessive calcification and bone formation.

The balance of RUNX2 and SOX9 regulates chondrogenesis. While RUNX2 controls the initial differentiation from BMSC to chondrocyte and the later hypertrophic growth and terminal differentiation, SOX9 is required for early chondrogenesis and the initial transition into hypertrophy. SOX9 expressing pre-hypertrophic chondrocytes express *Osx, MMP13, PTHrP* and begin to upregulate *RUNX2* to complete hypertrophy (Dy et al., 2012; Ikegami et al., 2011). RUNX2 regulation of chondrocyte maturation and hypertrophy requires it to suppress SOX9 activity in distinct stages of chondrocyte maturation (Cheng and Genever, 2010). Decreases in SOX9 expression are associated with longer hypertrophic zones and increased mineralization. Our data demonstrate that Kindlin-3 plays a role in the equilibrium between SOX9 and RUNX2. SOX9 blocks RUNX2 expression inhibiting hypertrophy and osteoblast differentiation in a dominant manner (Zhou et al., 2006). SOX9 upregulated *Bapx1/Nkx3.2* suppresses RUNX2 in chondrocytes preventing late-stage chondrogenesis (Yamashita et al., 2009; Provot et al., 2006). Thus, the increase in *SOX9* and decrease in *RUNX2* in Kindlin-3 deficient BMSCs and the loss of integrin binding in the Kindlin-3 mutant mice results in faster progression through the early stages of chondrocyte differentiation but slows terminal differentiation resulting in the retention of chondrocyte cells in the ossification front.

### Kindlin’s integrin binding function in the regulation of chondrogenesis

Downstream of SOX9 activation, chondrocytes begin to enhance their secretion of extracellular matrix proteins, requiring increased integrin activation and binding during the transition into hypertrophy (Quintana et al., 2009). The extreme changes in chondrocyte shape and ECM during chondrogenesis requires both inside-out and outside-in integrin signaling and activation. In the growth plate, chondrocytes undergo physical relocation and rotation requiring movement through the extracellular matrix, rearrangement of focal adhesions, and integrin activation. Chondrocyte adhesion and maturation is regulated by integrins, and integrin expression can be altered with maturation. Studies have suggested that integrin signaling, particularly through β_1_ integrin, may have a temporal role during chondrogenesis. For example, α_5_β_1_ integrin is expressed in the proliferative and hypertrophic zones (Enomoto-Iwamoto et al., 1997). The β_1_ integrin controls later stages of chondrogenesis as inhibition of the β_1_ integrin-induced the expression of SOX9 and collagen type X at early time points (Lu et al., 2008). Further, β_1_ integrin deletion in articular cartilage resulted in disorganized cartilage with disruption of the actin cytoskeleton and enhanced terminal differentiation (Raducanu et al., 2009). In our model of early chondrogenesis, we do not see a disruption of the actin cytoskeleton but rather an increase in actin formation. However the loss of integrin activation due to Kindlin-3 deficiency did accelerate differentiation. The effects are likely less pronounced since the other Kindlin isoforms are still able to perform their integrin binding function. In addition, the β_1_ integrin-deficient articular cartilage demonstrated a thicker zone of calcified cartilage as demonstrated by collagen type X staining (Raducanu et al., 2009), similar to the increased collagen type X expression seen with Kindlin-3 mutation. MMP expression was not altered by loss of β_1_ integrin expression in mesenchymal cells. Deletion of the β_1_ integrin in mesenchymal cells results in chondrocyte clustering and structural disorganization. Expression of collagen type X started earlier in the cartilage (Raducanu et al., 2009). Blocking integrin β_1_ in adipose-derived stem cells cultured in control media upregulated collagen type X and SOX9 gene expression (Lu et al., 2008) similar to our findings with Kindlin-3 deletion indicating that blocking integrin activation induces a more prominent chondrogenic phenotype prior to induction with chondrogenic media.

In the patient described here, Kindlin-3 deficiency significantly affected integrin activation and chondrocyte hypertrophy. Using a constitutively activated integrin rescue, we determined that of integrin activation could rescue chondrocyte adhesion and prevent the excessive *SOX9* expression. In concert, Kindlin-3 mutated growth plates in which integrin binding only was altered, chondrocyte hypertrophy was again affected and SOX9 expression enhanced. Taken together, these data demonstrated that the integrin binding of Kindlin-3 was responsible for the changes in chondrocyte differentiation. These data are in line with previous studies demonstrating that alterations in integrin binding proteins affected chondrogenesis. Deletion of integrin-linked kinase (ILK) results in shortened and disorganized growth plates due to decreased hypertrophic region due to abnormal chondrocyte proliferation and altered cell shape (Grashoff et al., 2003; Terpstra et al., 2003). Loss of ILK results in 30% reduced chondrocyte adhesion to fibronectin and collagen type I (Grashoff et al., 2003) and ∼50% to collagen type II (Terpstra et al., 2003). However, unlike the increased actin fiber formation seen in our study, ILK-deficient chondrocytes demonstrated fewer, shorter, and disorganized stress fibers (Grashoff et al., 2003). Further, proliferation was decreased in the ILK null chondrocytes *in vivo*. These discrepancies are partially due to studying the BMSCs prior to chondrogenic differentiation. Similar to our findings with Kindlin-3, expression of another integrin binding protein, vinculin in increased with chondrocyte maturation. Vinculin deletion in chondrogenic mesenchymal cells suppresses RUNX2, collagen type II, collagen type X, and aggrecan, but does not affect SOX9 expression. In addition, loss of vinculin resulted in disrupted columnar proliferation and decreased hypertrophy (Koshimizu et al., 2012).

Maturation of chondrocytes is associated with changes in the actin filament structure with chondrocyte dedifferentiation being associated with actin disruption (Benya et al., 1978; Zanetti and Solursh, 1984; Hirsch et al., 1996; Brown and Benya, 1988). Thus, changes in the actin cytoskeleton of BMSCs in monolayer may be representative of changes in chondrogenic differentiation similar to those seen with primary chondrocytes (Kerr et al., 2008). Disruptions in the actin cytoskeleton can also control chondrocyte maturation (Zanetti and Solursh, 1984; Kerr et al., 2008). MSCs treated with cytochalasin D to disrupt the actin network spontaneously undergo chondrogenesis. Thus, the diminished cell-matrix interactions caused by Kindlin-3 deficiency may promote a more chondrogenic phenotype in BMSCs. Kindlin-2 knockdown in pre-osteoblast cells resulted in decreased proliferation and spreading due to decreased activation of Rac1. However integrin expression and activation were unchanged (Jung et al., 2011). Correspondingly, Kindlin-3 regulates Rac1 activation downstream of integrin activation (Xue et al., 2013). Kindlin-3 regulation of integrin activation is required for cytoskeletal rearrangement. Kindlin deficient cells, platelets, in particular, have reduced spreading on a variety of substrates. One study demonstrated that Syk and Vav1 signaling downstream of integrin activation and Kindlin3 regulated Rac1 and Cdc42 to control cell spreading (Xue et al., 2013) Rac1 activation and subsequent reorganization of the actin cytoskeleton plays an important role in the transition to hypertrophy (Kerr et al., 2008).

### Kindlins alter bone development beyond their roles in chondrogenesis

A recent report of a new IADD (LAD-III) patient demonstrated that a loss of Kindlin-3 resulted in increased bone density with thickened cortices and trabeculae. Bars of calcified cartilage remained within the mature bone, and the hypertrophic cartilage was expanded with increased sclerotic areas. The cartilage below the proliferative zone was disorganized (Crazzolara et al., 2015). After BMT, the cartilage contained more mature cartilage tissue and re-established columnar organization. A recent biopsy of an iliac crest from a patient with a Kindlin-3 mutation demonstrated abnormal osteoclasts numbers as well as cartilage remnants in the trabecular bone (Palagano et al., 2017). Another patient with a Kindlin-3 mutation displayed similar remnants of calcified cartilage in the trabecular bone as well as sclerotic primary spongiosa in the subchondral region (Crazzolara et al., 2015), which is similar to that seen in the K3KI mutant mice.

Our data demonstrate that Kindlin-3 and Kindlin-2 play opposing roles in the regulation of bone development. Kindlin-2 expression in MSCs during chondrogenesis was required for chondrocyte proliferation and development of the primary ossification center (Wu et al., 2015). We demonstrate that Kindlin-3 loss increased MSC proliferation and that Kindlin-3 was not expressed in proliferative chondrocytes in growth plates. Contrary to Kindlin-2, Kindlin-3 regulated chondrocyte hypertrophy. The two Kindlin proteins also interacted differently with the transcription factors associated with chondrocyte differentiation. The Kindlin-2 loss reduced SOX9 expression, while Kindlin-3 loss induced SOX9 expression in MSCs. Similar opposing effects were seen with collagen type II mRNA. These data indicated Kindlin-2 partners with SOX9 expression to drive early chondrocyte differentiation, while Kindlin-3 and RUNX2 regulate chondrocyte hypertrophy. Thus, Kindlin-3 and Kindlin-2 are expressed distinctly in MSCs and chondrocytes, and both orthologs are required for chondrogenesis to proceed normally.

There is also a potential role for Kindlins in osteogenesis since integrin activation is also integral to osteoblast differentiation. For example, α_5_ integrin promotes BMSC differentiation down the osteoblast lineage (Hamidouche et al., 2009). Overexpression or activation of α_5_ integrin-induced RUNX2, alkaline phosphatase, and collagen type I expression which controls the osteogenic lineage. Thus, an interaction between RUNX2 and integrin activation may control bone development. Inactivation of β_1_ integrin in osteoblasts results in decreased bone mass (Zimmerman et al., 2000). In addition, the expression of integrins, including α_5_β_1_, is required to drive osteoblast differentiation from human MSC (Hamidouche et al., 2009; Moursi et al., 1997). Deficiency of migfilin, a Kindlin binding protein, also results in severe osteopenia (Xiao et al., 2012; Brahme et al., 2013). Similar to our results, loss of migfilin resulted in decreased adhesion of MSCs (Brahme et al., 2013). Deletion of migfilin (filamin-binding LIM protein-1), another protein controlling integrin activation that binds to Kindlins, also induces osteopenia in mice. Similar to our data with the loss of Kindlin-3, migfilin deficiency in BMSCs lead to decreased cell adhesion to collagen type I and fibronectin (Xiao et al., 2012). BMSC differentiation into osteo-chondroprogenitor cells then chondrocytes. In addition to osteogenesis, Kindlin-3 has a known role in osteoclastogenesis. While, initial studies of Kindlin-3 in bone concentrated on the effect of Kindlin-3 in osteoclast function (Schmidt et al., 2011); their data also showed significantly decreased bone formation rate to bone surface area in Kindlin-3 deficient animals. Additionally, Kindlin-3 deficiency in patients alters the hematopoietic niche. Kindlin-3 expression in hematopoietic stem cells is required for their continued maintenance in the bone marrow (Ruppert et al., 2015). The inability of Kindlin-3 to bind integrins in our mice may result in alterations in crosstalk between MSCs and hematopoietic stem cells.

In summary, we demonstrate that Kindlin-3 plays an important role in the differentiation of mesenchymal stem cells down the chondrocyte lineage. This represents a potential secondary mechanism leading to osteopenia in Kindlin-3 deficient patients. Additional studies will need to be completed to understand the relationship between RUNX2, SOX9, and Kindlin-3 during chondrogenesis.

## AUTHOR CONTRIBUTIONS

Conceptualization: B.A.K., T.V.B.

Formal Analysis: B.A.K., L.S., A.H.J.

Funding Acquisition: T.V.B.

Investigation: B.A.K., L.S., A.H.J., D.P.L.

Project Administration: B.A.K, T.V.B

Resources: J.S.W., D.P.L., A.I.C.

Supervision: B.A.K., T.V.B., A.I.C.

Visualization: B.A.K., L.S.

Writing – Original Draft: B.A.K., T.V.B.

Writing – Review and Editing: B.A.K., L.S., A.H.J., J.S.W., D.P.L., A.I.C., T.V.B.

## The abbreviations used are

3D: three dimensional
BMSC: Bone marrow-derived mesenchymal stem cell
BSA: Bovine serum albumin
DMEM-LG: Dulbecco’s Modified Eagle’s Medium-Low Glucose
IADD: integrin adhesions deficiency disease
K3KI: Kindlin-3 knock-in
LAD: leukocyte adhesion deficiency
MSC: mesenchymal stem cell
TGF: transforming growth factor
TRAP: tartrate-resistant acid phosphatase

## ACKNOWLEDGEMENTS

We acknowledge funding from NIH/NHLBI (HL073311 and HL071625) to Dr. Byzova. Dr. Kerr was supported by a Ruth L. Kirschstein NRSA award (F32 CA142133) followed by a Pathway to Independence Award (K99 CA175291) from the NIH/NCI. We thank Joanna Ireland and Dr. Finke from the Department of Immunology for the gift of several FACS antibodies. We thank Dr. Sara Tomechko for assistance with the LiCor imaging and reagents and Dr. Young-Woong Kim for assistance with primer design. We thank Dr. Plow and his lab members for the gifts of fibrinogen, for the Kindlin-3 antibodies, and for their technical assistance. We thank Brandi Bickford of the Virtual Microscopy Core for her help with slide scanning. The authors declare no competing financial conflicts of interest.

