## Supplemental Figures for "Kindlin-3 Mutation in Mesenchymal Stem Cells Results in Enhanced Chondrogenesis"

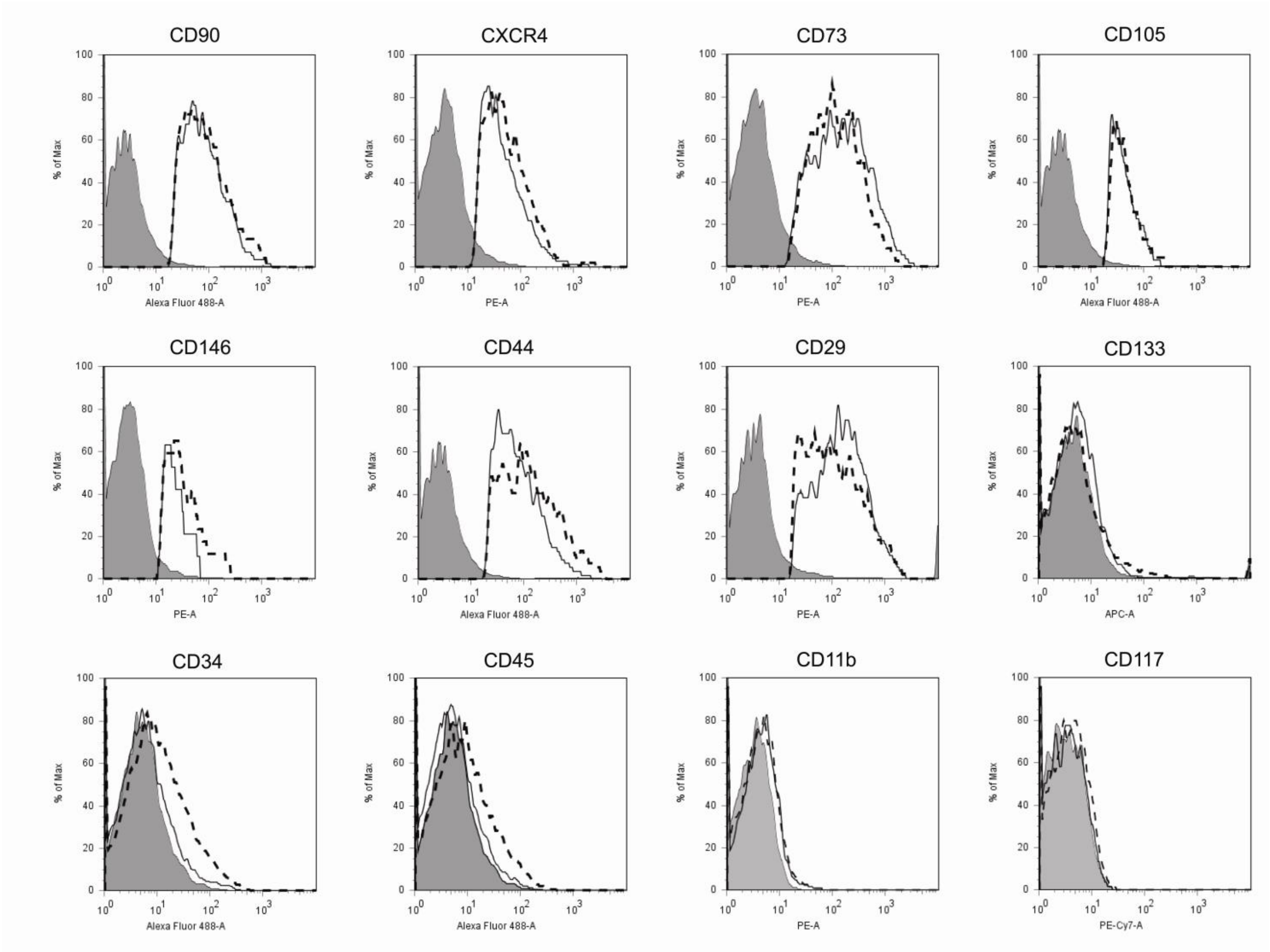


**Figure S1. Characterization of BMSC Markers**. Normal (solid line) and subject (dotted line) BMSCs were analyzed for MSC and HSC markers and compared to isotype controls (gray filled).

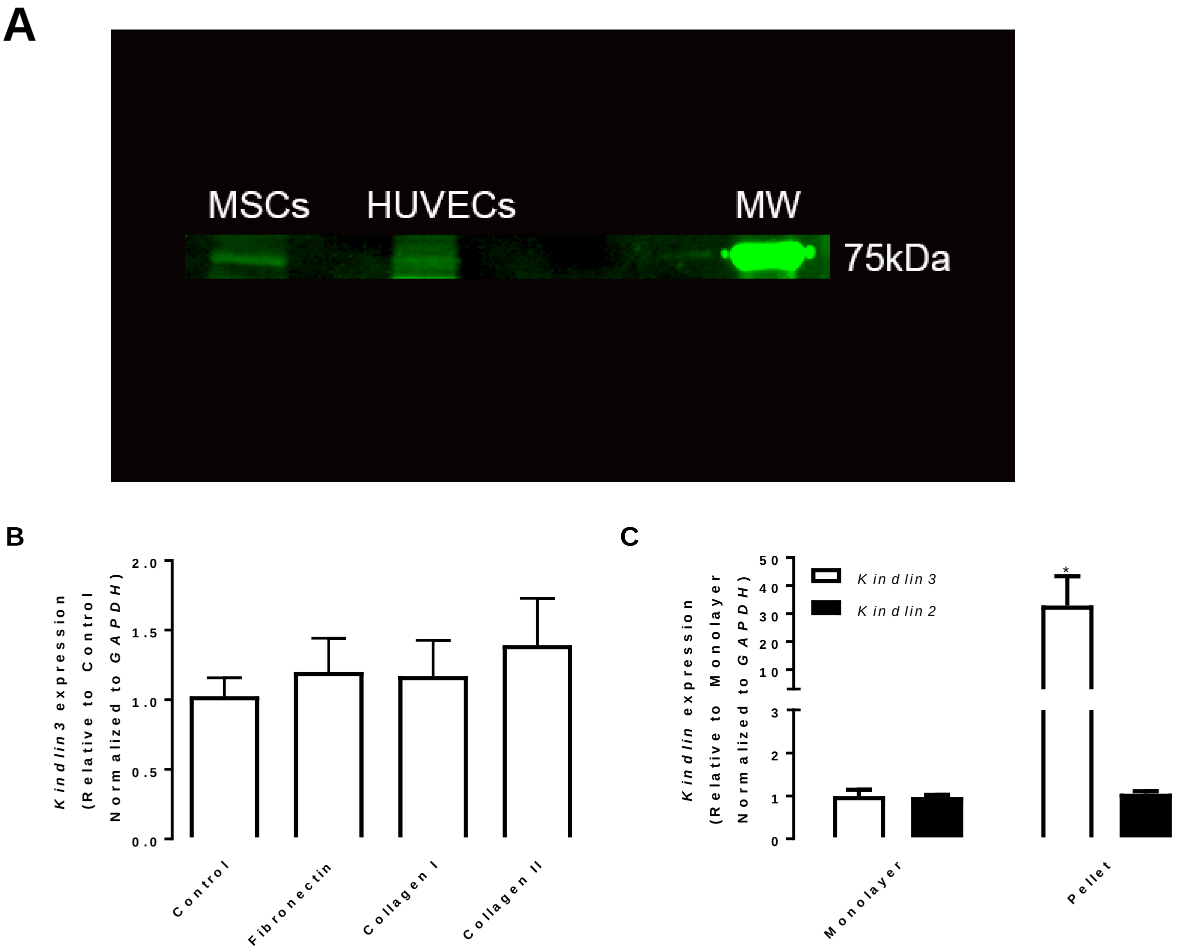


**Figure S2. Kindlin3 Expression in Normal BMSCs.** BMSCs were isolated from normal patients and analyzed for Kindlin-3 expression. **A**, BMSCs and human vein endothelial cells (HUVECs) were lysed and immunoblotted for Kindlin-3. **B**, BMSCs were plated on tissue culture plastic (Control), fibronectin, collagen type I, or collagen type II. represented as mean fold change from control±SEM. **C**, BMSCs were plated in monolayer or allowed for form 3D pellets. Gene expression of *Kindlin2* and *Kindlin3* were measured relative to *GAPDH* represented as mean fold change from monolayer±SEM. * represents $p<0.05$ by one-way ANOVA.

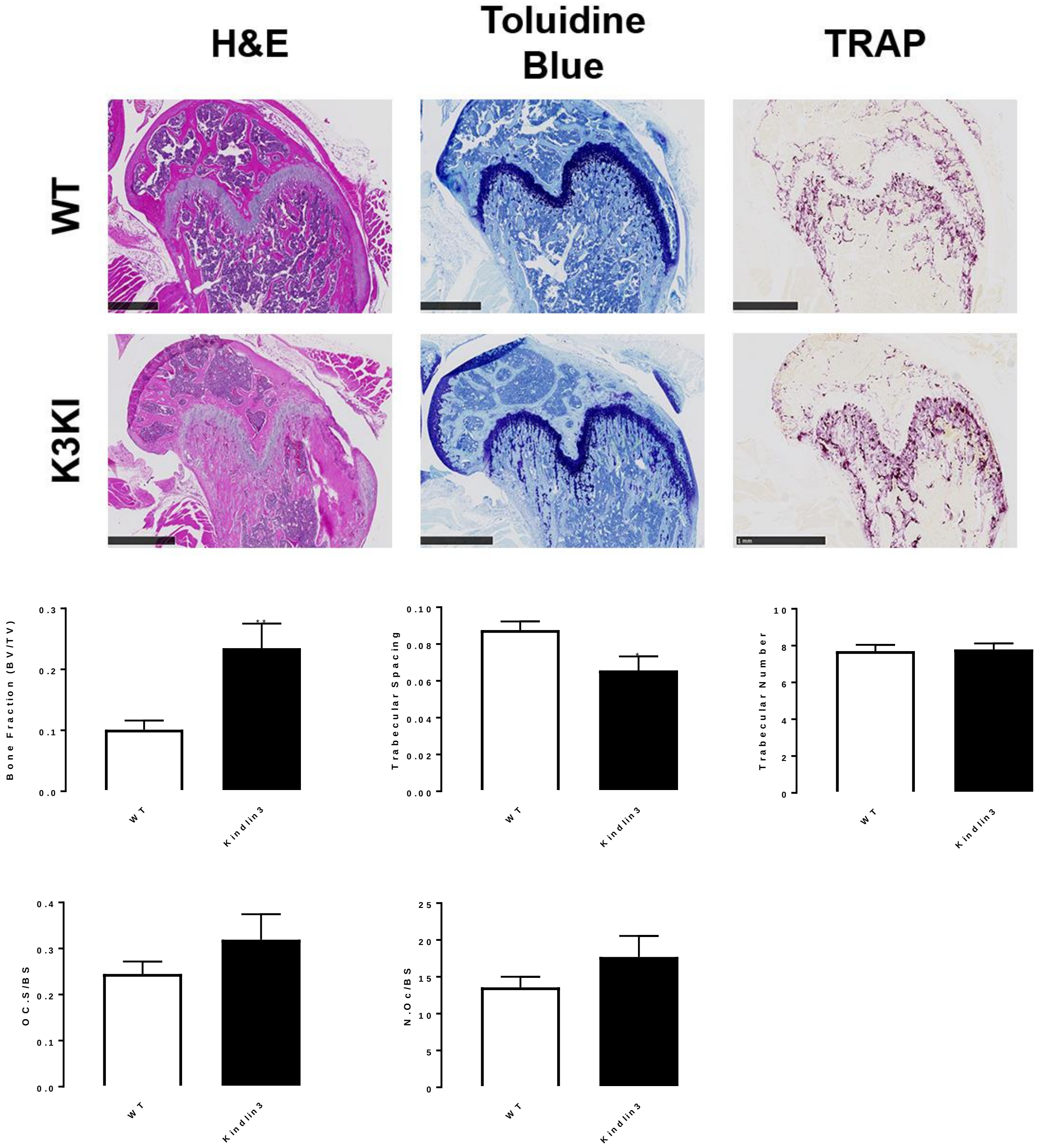


**Figure S3. Bone histomorphometry and osteoclast numbers in K3KI bones.** Bones collected from 9-week old wildtype (WT) and K3KI mice were decalcified, sectioned, and stained with H&E or TRAP. Bone histomorphometry and osteoclast numbers were calculated using Bioquant Osteo. Scale bars represent 1mm. * represents $p<0.05$ and ** represents $p<0.01$ by Student's *t* test.
